# Cortical and thalamic influences on striatal involvement in instructed, serial reversal learning; implications for the organisation of flexible behaviour

**DOI:** 10.1101/2021.12.20.472804

**Authors:** Brendan Williams, Anastasia Christakou

## Abstract

Cognitive flexibility is essential for enabling an individual to respond adaptively to changes in their environment. Evidence from human and animal research suggests that the control of cognitive flexibility is dependent on an array of neural architecture. Cortico-basal ganglia circuits have long been implicated in cognitive flexibility. In particular, the role of the striatum is pivotal, acting as an integrative hub for inputs from the prefrontal cortex and thalamus, and modulation by dopamine and acetylcholine. Striatal cholinergic modulation has been implicated in the flexible control of behaviour, driven by input from the centromedian-parafascicular nuclei of the thalamus. However, the role of this system in humans is not clearly defined as much of the current literature is based on animal work. Here, we aim to investigate the roles corticostriatal and thalamostriatal connectivity in serial reversal learning. Functional connectivity between the left centromedian-parafascicular nuclei and the associative dorsal striatum was significantly increased for negative feedback compared to positive feedback. Similar differences in functional connectivity were observed for the right lateral orbitofrontal cortex, but these were localised to when participants switched to using an alternate response strategy following reversal. These findings suggest that connectivity between the centromedian-parafascicular nuclei and the striatum may be used to generally identify potential changes in context based on negative outcomes, and the effect of this signal on striatal output may be influenced by connectivity between the lateral orbitofrontal cortex and the striatum.

## Introduction

Flexible control over behaviour enables goal directed action by allowing an individual to respond to changes in its environment. Being adaptive to change means an individual can adjust their behaviour while maintaining their current goal, which is particularly important in ambiguous or volatile environments where there is uncertainty around optimal behaviours. One common behavioural assay is the reversal learning task (Izquierdo et al., 2017; Yaple & Yu, 2019). During reversal learning participants learn associations between actions and outcomes, with the aim of maximising reward. Some actions are more rewarding, while others are less so. These associations between actions and outcomes change during the task and therefore participants need to alter how they respond to continue maximising reward. Perseverative responding, the inability to effectively switch to an alternate response strategy following reversal, is indicative of inflexibility, and elevated in conditions characterised by repetitive behaviour including addiction (De Ruiter et al., 2009; Ersche et al., 2011; Verdejo-Garcia et al., 2015), autism (Crawley et al., 2020; though also see D’Cruz et al., 2016) and obsessive compulsive disorder (Remijnse et al., 2006).

Cognitive flexibility requires the orchestration of signals from across the cortical surface and subcortical brain regions. Previous work has shown that the striatum, with its afferent projections from the cerebral cortex, thalamus, amygdala, and hippocampus (among other regions) is well positioned to act as an integrative information hub (Haber, 2016) and supports reversal learning (Bradfield, Hart, et al., 2013; Cools et al., 2002; Ruge & Wolfensteller, 2016). One striatal afferent consistently implicated in studies of reversal learning is the orbitofrontal cortex (Dalton et al., 2016; Rudebeck & Murray, 2008; Rygula et al., 2010; Tsuchida et al., 2010). Reduced orbitofrontal activation during reversal learning is associated with increased inflexibility in obsessive compulsive disorder, (Remijnse et al., 2006), and with a reduced capacity to use negative outcomes to guide behaviour (Groman et al., 2019). Within the context of reversal learning, the orbitofrontal cortex is broadly thought to be involved in the shifting of behaviour following the reversal of outcome contingencies (Izquierdo et al., 2017; Uddin, 2021). More specifically, the medial and lateral portions of the orbitofrontal cortex are thought to have dissociable roles, with the former involved in outcome evaluation, and the latter representing the current state of the task and its associated contingencies (Hampshire et al., 2012; Hervig et al., 2020; Noonan et al., 2017). Therefore, though the medial division of the orbitofrontal cortex is generally important for goal directed learning, the lateral portion is specifically involved in reversal learning and changing behaviour following the reversal of reward contingencies (Dalton et al., 2016; Hampshire et al., 2012; Morris et al., 2016). Previous work from Bell, Langdon, et al. (2019) used a multi-alternative reversal learning task with a single uninstructed reversal to investigate the neural mechanisms underlying reversal learning. This task closely mimics the setup of animal studies of reversal learning. It has a protracted period of initial and reversal learning where participants learn correct and incorrect choices from experiencing trial-and-error. These protracted learning periods are like learning in animal reversal learning task (e.g. in T-mazes Izquierdo et al, (2017)), where many trials are required for learning stimulus-outcome contingencies. This is because participants are not instructed about the reversal of reward contingencies. Instead, they rely on ongoing outcome processing to signal a potential change in environmental contingencies. Increased functional connectivity between the medial orbitofrontal cortex and ventral striatum was found after initial learning, and before reversal; at this time participants should have a stable initial representation of the task to guide goal-directed behaviour. This finding supports the proposed role of the medial orbitofrontal cortex in outcome evaluation and goal directed behaviour. Conversely, increased functional connectivity between the lateral orbitofrontal cortex and the dorsal striatum was found during reversal learning. The strength of functional connectivity was inversely correlated with trials to criterion during reversal learning and was found to be mediated by the learning rate for positive prediction errors. This suggests lateral orbitofrontal connectivity with the dorsal striatum promotes changes in behaviour following reversal via learning from positive outcomes, reducing response perseveration.

Another region important for cognitive flexibility is the thalamus. While traditionally viewed as a relay for sensory and motor signals, there is an increasing appreciation for the role of the thalamus in cognitive processes (Wolff & Vann, 2019). This includes cognitive flexibility and reversal learning, where several thalamic nuclei are known to have an important role. For instance, animal literature has shown that lesions to the mediodorsal nucleus of the thalamus does not impair initial discrimination learning but does impair reversal learning (Chudasama et al., 2001). More specifically, lesions to the mediodorsal thalamus are thought to impair reversal learning by preventing the use of recent history to guide future choice, and thus choice behaviour is increasingly stochastic following the reversal of reward contingencies (Chakraborty et al., 2016). In addition to the mediodorsal thalamus, the centromedian-parafascicular nuclei are also important for flexibility. However, while the mediodorsal projects primarily to the cortex with axon collaterals to the striatum, the centromedian-parafascicular nuclei project primarily to the striatum, with afferents that are preferentially connected with the striatal cholinergic system (Smith et al., 2009). Thalamic connectivity between the parafascicular nucleus of the thalamus (the rodent homologue of the primate centromedian-parafascicular (Smith et al., 2011)) and the striatal cholinergic system is important for reversal learning (Bradfield, Hart, et al., 2013); disconnection of these circuits impairs reversal learning by reducing striatal acetylcholine efflux, which in turn increases interference between new and existing contingency encoding following reversal (Bradfield, Bertran-Gonzalez, et al., 2013; Brown et al., 2010). Therefore, though both the mediodorsal and centromedian-parafascicular nuclei support cognitive flexibility, they show distinct roles and connectivity patterns.

Thalamostriatal connections between the centromedian-parafascicular nuclei and the striatal cholinergic system are also thought to be important for flexibility in humans. For instance, magnetic resonance spectroscopy has previously been used during uninstructed reversal learning, to demonstrate functionally relevant changes in dorsal striatal choline that are specific to the reversal of reward contingencies and are not present during initial learning (Bell et al., 2018). Moreover, baseline striatal choline levels are also associated with reversal learning performance, and suggests that the state of the striatal cholinergic system explains some variability in cognitive flexibility (Bell, Lindner, et al., 2019). Additionally, changes in functional connectivity between the centromedian-parafascicular nucleus in the thalamus and the dorsal striatum have been seen during reversal, but not initial learning in the same task (Bell, Langdon, et al., 2019). Based on evidence from animal and human literature we think the thalamostriatal connectivity between the centromedian-parafascicular nuclei and the dorsal striatal system is important for producing internal representations that are context dependent and support cognitive flexibility. More specifically, thalamostriatal connections are believed to recruit cholinergic interneurons to support new learning without “overwriting” prior knowledge (Bradfield, Bertran-Gonzalez, et al., 2013). Cholinergic involvement, alongside dopaminergic prediction errors, enables behaviour that can adaptively and efficiently respond to change by making the system sensitive to the broader behavioural context, beyond simply outcome contingencies. Furthermore, the striatal cholinergic system may enable the representation of context by modulating the output of striatal projection neurons (Stayte et al., 2021). Therefore, the concurrent representation of context-dependent contingencies in the striatum facilitates flexible behaviour that is responsive to change. Indeed, this preposition is supported by computational work from Franklin & Frank (2015), who show the inclusion of cholinergic interneurons in a model of the basal ganglia enables learning rates to respond to environmental noise and uncertainty. Cholinergic interneurons are tonically active and show a transient pause and rebound response to phasic activity. Changes in length of this transient pause in Franklin & Frank’s (2015) model were associated with balancing flexibility and stability, with pause length inversely associated with learning rate during reversal. Shorter pauses facilitated faster updating of expected values but meant the system was more perturbed by noise, while the inverse was true for longer pauses. Importantly, a pause length that was reciprocally modulated by medium spiny neuron activity (compared to models with fixed pause lengths), enabled the system to respond adaptively to change, minimising errors during probabilistic reversal. These dynamics may ultimately regulate the effects of dopamine on the plasticity of corticostriatal synapses, enabling the striatum to generate internal representations that are responsive to change and uncertainty in the environment. Furthermore, these dynamics are in line with our understanding of how variability in cholinergic transmission influences sensitivity to volatility and noise during reversal learning.Bell, Langdon, et al., 2019; Bell, Lindner, et al., 2019), we aimed to bridge the gap between animal and human work. We used a task homologous with animal studies to investigate how cortical, striatal, and thalamic systems interact during initial and reversal learning in humans. However, most reversal learning tasks used in human neuroimaging work are more like serial reversal learning than the multi-alternative task of the Bell et al. studies. Internal representations of task context are readily acquired in serial reversal learning, and once an “if not A, then B” heuristic for correct and incorrect choices exists, no additional information is relevant for representing task structure. Therefore, task representation in serial reversal learning may be considered as “saturated”. However, task representations for the multi-alternative task can be considered as “unsaturated”, since participants are not instructed on the structure of the task and only compile mature task representations following both the protracted initial and reversal learning periods. Therefore, these differences in task representation may explain evidence suggesting dissociable roles for choline in serial and multi-alternative reversal learning (Bell, Lindner, et al., 2019; Williams & Christakou, 2021). We propose having a higher cholinergic “tone” at rest is beneficial under a saturated task representation, by limiting learning disproportionately from probabilistic and regressive errors, thereby promoting stability. By contrast, a lower tone is beneficial for unsaturated task representations, by maximising contrast between periods of stability and change, thereby promoting flexibility.

It is unclear how cortical and thalamic regions interact with the striatum and produce flexible behaviour in a context with a saturated task representation. In such a context, there is no new contextual information following initial discrimination and reversal learning, because the reversal has been instructed. Therefore, connectivity specifically from the centromedian-parafascicular nuclei to the striatum may not contribute to serial reversal learning in the same way as in the uninstructed multi-alternative task, where the reversal needs to be discovered. Instead, behaviour may be mediated via cortically driven mechanisms, using prior knowledge to guide action in line with the known task representation. This is supported by evidence demonstrating that inputs from the orbitofrontal cortex to striatal cholinergic interneurons are necessary for generating internal representations in the striatum (Stalnaker et al., 2016). Alternatively, these thalamic connections may continue to be important after initial and reversal learning. For instance, they signal behaviourally relevant sensory events (Matsumoto et al., 2001; Schepers et al., 2017), and respond to changes in context (Yamanaka et al., 2018). Therefore, thalamostriatal connectivity may be relevant for identifying contextual change to promote flexibility in serial reversal as in the multi-alternative task.

To arbitrate between these possibilities, we use recent advances in parcellation approaches and functional magnetic resonance acquisition optimised for spatial specificity to examine the roles of orbitofrontal, striatal, and thalamic regions in producing cognitive flexibility in serial probabilistic reversal learning (Iglesias et al., 2018; Volz et al., 2019). In our task we define two distinct phases that appear consistently over successive reversal episodes. The “*re-learning phase*” is the period between the reversal of outcome contingencies and participants reaching the learning criterion. Once they reach criterion participants are in the “*stability phase*” until outcome contingencies reverse again. We used psychophysiological interaction analysis to study the functional connectivity between regions of interest in subdivisions of the cortex, striatum, and thalamus during re-learning and stability. Additionally, we conducted exploratory analyses investigating how reward and error signals may be used to guide behaviour during re-learning and stability phases.

## Methods

### Participants

36 healthy adult participants (mean age = 24.38 years; SD = 6.51; range = 18-49; 23 female) were recruited via opportunity sampling from the University of Reading community. Participants were eligible for participation if they were right-handed, and did not self-report the use of cigarettes, recreational drugs, or psychoactive medication or have a formal diagnosis of psychiatric or mental health condition. Participants were reimbursed £15 for their time. Seven participants were excluded from the final dataset. One participant had artifacts in MR data due to braces; one participant had technical errors during scanning; one participant had registration issues; four participants responded on less than 95% of trials during the task.

### Materials

Magnetic resonance images were collected on a Siemens MAGNETOM Prisma 3T MRI Scanner with a 32-channel receiver head coil at the Centre for Integrative Neuroscience and Neurodynamics at the University of Reading. The probabilistic reversal learning task was programmed using MATLAB (2017b, The Mathworks, Inc, Natick, MA, United States) and Psychtoolbox-3 (Brainard, 1997) on a Macintosh running macOS Sierra. The task was presented on a BOLDscreen LCD (Cambridge Research Systems Ltd, Rochester, Kent, United Kingdom) during scanning and displayed to the participant via a mirror placed above their eyes. Task presentation, synchronised with functional volume acquisition, was controlled by a computer running Windows 7, MATLAB 2015b and Psychtoolbox-3.

### Learning task

Two abstract images of fractal patterns were shown on the left and right hemifield of the visual display. Participants had to choose one of the two images within 2000ms by pressing the corresponding button on a button box, else a “too late” message was displayed. After selection, the chosen stimulus was immediately highlighted (duration: min = 1500ms, max = 2500ms, 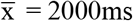, plus a jitter sampled from an exponential distribution (min = 500ms, max = 6000ms, 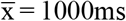)). The outcome of the participant’s choice was then presented for 1000ms. A jittered fixation cross sampled from an exponential distribution (min = 500ms, max = 3000ms, 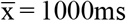) followed and was presented in the centre of the display. The participant’s cumulative points total was then presented for 500ms and the trial ended with a second jittered fixation cross in the centre of the screen, sampled from a different exponential distribution to the first fixation cross (min = 500ms, max = 6000ms, 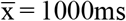). Delays and jitters were optimised to separate events of interest without compromising the psychological features of the trial, desynchronise trial timings from the fMRI acquisition repetition time, and to fully sample across the hemodynamic response function. Figure 1 shows a schematic of the task trial structure and timings.

**Figure 1:**
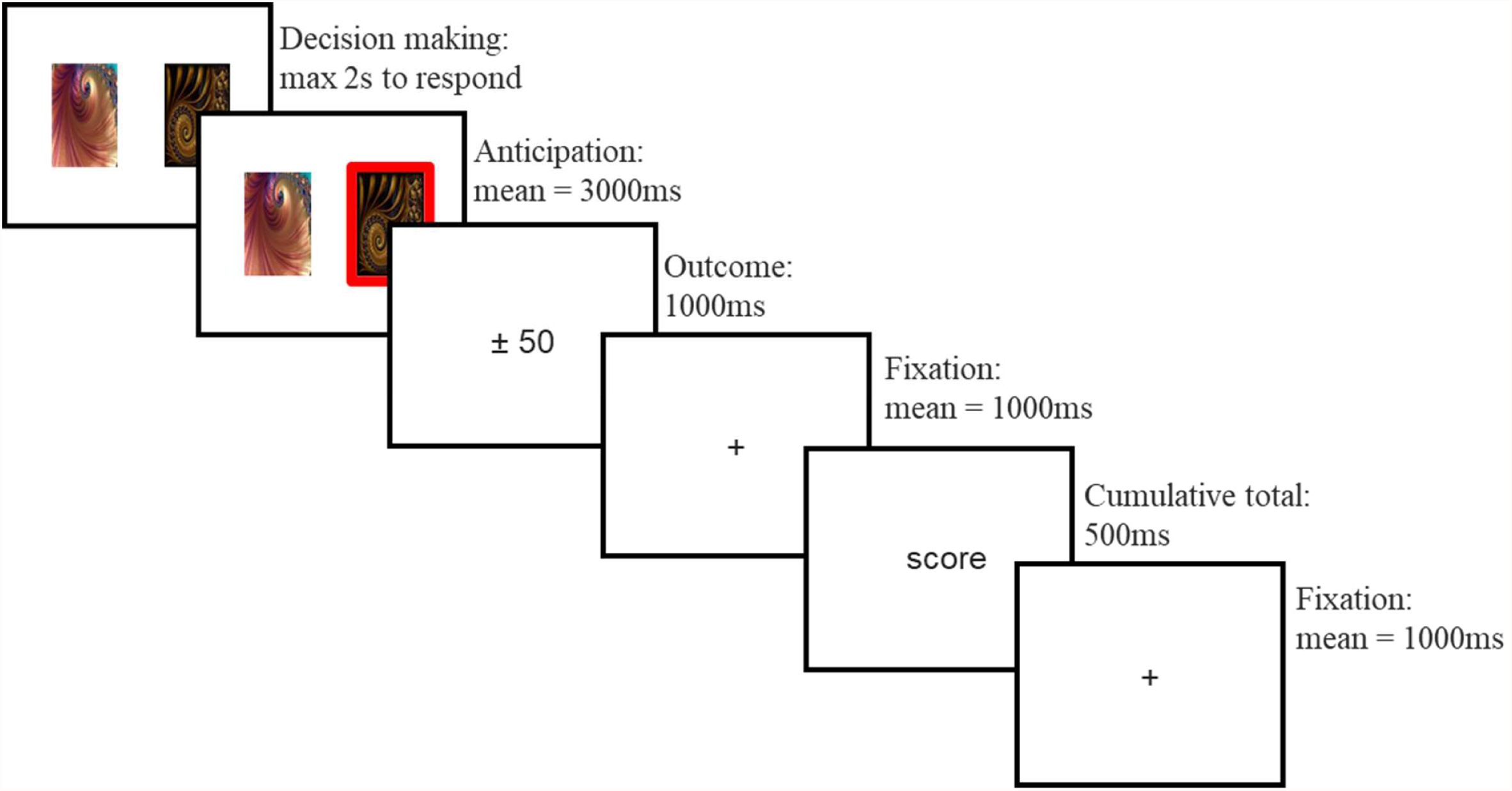
Overview of a single trial. Participants are initially shown two abstract fractal images and given two seconds to choose one image. Their choice is then highlighted. The participant is then shown the outcome of their choice; this will either be an increase or decrease of 50 points if they selected an image, or 0 points if they made no choice. The outcome is followed by a fixation cross, their cumulative total so far, and finally another fixation cross.

At the beginning of the task, one of the two images were randomly assigned as the correct image, and the other as the incorrect image. The probability of winning points on the correct image was 0.8, and the probability of losing points was 0.2. The inverse was true for the incorrect image. Outcomes were pseudo-randomised such that the assigned probabilities were true for blocks of 20 consecutive selections of the correct or incorrect choice. Additionally, no more than six of the same outcomes (win or loss) would be consecutively presented for the correct or incorrect choice. If participants won, their cumulative total increased by 50. If they lost, their cumulative total decreased by 50. If they did not choose an image, their cumulative total did not change. For outcome probabilities to reverse participants had to reach and maintain a predefined learning criterion: the selection of the correct image on five of the previous six trials. After reaching criterion participants entered a *stability phase* where the probability of reversal was equal to the number of trials where criterion had been maintained, divided by 10 (adapted from (Hampton et al., 2006)). If criterion was not maintained, then the probability of reversal was reset to 0 and restarted once criterion was reached. This variable length was included to minimise identification of the learning criterion and anticipation of the reversal event. The reversal event involved the switching of outcome probabilities, with the correct image becoming incorrect and *vice versa*. On reversal, participants had to re-reach and maintain the learning criterion for the reassigned outcome probabilities (*re-learning phase*) before outcome probabilities would reverse again. Participants completed 360 trials of the reversal learning task. There was no limit on the number of reversal events a participant could experience. Left-right stimulus presentation was randomised across trials. Before starting, participants were instructed that: their aim was to collect as many points as possible; the outcome was dependent on the image they chose; one choice may be better than the other; and the better choice could change during the task. Participants completed 20 practice trials prior to entering the MRI scanner. Practice trials followed the same structure as trials in the scanner, but participants did not receive any feedback for their choices. Instead, hashtags were presented in place of outcome and cumulative total feedback.

### fMRI acquisition

T2-weighted whole-brain blood oxygen level-dependent (BOLD) functional images were acquired using a multi-band 2D-echo-planar imaging sequence with GeneRalized Autocalibrating Partially Parallel Acquisitions (GRAPPA) (acceleration factor = 2) [TR = 2160ms; TE = 30ms; slices = 93; voxel volume ≈ 1.6mm^3^; slice thickness = 1.6mm; distance factor = 0%; FOV = 205 × 205 mm; matrix = 128 × 128; flip angle = 90°; multiband acceleration factor = 3; phase encoding direction = A → P (negative polarity); interleaved acquisition; echo spacing = 0.75ms; fat suppression]. BOLD acquisition parameters were optimised for spatial resolution due to the scale of thalamic nuclei relative to standard parameters used at 3T (2.5-3mm isotropic), and because smaller voxels decreases signal loss in the orbitofrontal cortex as field inhomogeneities cause partial volume distortions for fewer voxels (Volz et al., 2019; Weiskopf et al., 2007). Fieldmap images were acquired to correct for distortions in the acquired data due to inhomogeneities in the magnetic field [TR = 2900ms; TE = 53.8ms; slices = 93; voxel volume ≈ 1.6mm^3^; slice thickness = 1.6mm; distance factor = 0%; FOV = 205 × 205 mm; matrix = 128 × 128; flip angle = 90°; multiband acceleration factor = 3; phase encoding direction = A → P and P → A; interleaved acquisition; echo spacing = 0.75ms; fat suppression; GRAPPA acceleration factor = 2]. High resolution T1-weighted anatomical images were acquired with a magnetization-prepared rapid gradient-echo (MP-RAGE) sequence with GRAPPA (acceleration factor = 2) [TR = 2300ms; TE = 2.29ms; TI = 900ms slices = 192; voxel volume ≈ 0.9mm^3^; slice thickness = 0.94mm; distance factor = 50%; slice oversampling = 16.7%; FOV = 240 × 240mm; matrix = 256 × 256; flip angle = 8 °; phase encoding direction = A → P; echo spacing = 7ms].

### Analysis of fMRI data

fMRI image analysis was performed principally using the FSL (6.0.4) toolbox from the Oxford Centre for Functional MRI of the Brain (FMRIB’s Software Library, www.fmrib.ox.ac.uk/fsl), and the FreeSurfer image analysis suite (version 6.0.0).

### Preprocessing

fMRI data pre-processing was carried out using FEAT (FMRI Expert Analysis Tool) Version 6.00. Registration of high resolution structural, functional and standard space images was carried out using FLIRT (Normal search, 12 DOF) (Jenkinson et al., 2002; Jenkinson & Smith, 2001); structural registration to standard space was further refined using FNIRT nonlinear registration (Normal search, 12 DOF, warp resolution = 10mm) (Andersson et al., 2007a, 2007b). MCFLIRT was used to identify motion artefacts in functional data. Motion and distortion correction were simultaneously applied to functional data using MCFLIRT parameters and B0 unwarping parameters from fieldmap images respectively. Functional data were spatially smoothed using a smoothing kernel (3.2 FWHM).

### Anatomical segmentation

Cortical reconstruction and volumetric segmentation were performed with FreeSurfer. Firstly, anatomical images were processed using the FreeSurfer recon-all pipeline, which included the generation of subject specific parcellations of the medial and lateral orbitofrontal cortex. We then used the ThalamicNuclei tool in FreeSurfer to perform thalamus segmentation. This tool performs segmentation of the thalamus using Bayesian Inference, and is based on a probabilistic atlas constructed from histological and *ex vivo* MRI data (Iglesias et al., 2018). Subject space masks for the centromedian, parafascicular, mediodorsal (medial), and mediodorsal (lateral) nuclei of the thalamus and the medial and lateral portions of the orbitofrontal cortex were generated from the automated FreeSurfer parcellations. Masks were transposed from anatomical space to functional space using FLIRT affine transformation parameters and re-binarised. Centromedian and parafascicular nuclei masks were combined to create a single mask for the Centromedian-parafascicular complex; mediodorsal (medial), and mediodorsal (lateral) masks were combined to create a mask for the mediodorsal nucleus.

### Statistical analysis

First level and higher-level statistical analyses for the fMRI data were performed using FEAT. At the first level, two general linear models were run.

#### Model 1 – trial-wide modelling for activation comparison across task phases

The first modelled different trial types as separate regressors. These regressors were modelled as boxcar functions, the onset coincided with the onset of choice stimuli (the start of the trial as depicted in Figure 1) and the offset with the removal of the cumulative total (the start of the intertrial fixation cross; Figure 1). Twelve regressors were used to model different trial types at the subject level; Trial regressors are defined in Table 1, and are based on the phase of the task (initial learning, re-learning, or stability), the choice made (correct or incorrect), and the outcome of the choice (positive or negative feedback). A regressor to separate final reversal errors (Figure 2) from other reversal errors was also included. Each regressor and its temporal derivative were convolved with the canonical (double gamma) haemodynamic response function. Activation maps for each regressor were generated at the subject level by creating contrast estimates relative to an implicit baseline derived from fixation periods (Figure 1). Contrast estimates for this model were generated at the whole-brain level.

**Table 1.** Definitions of trial types within the task used for the analysis of functional magnetic resonance imaging data based on the task phase, the participant’s choice, and the outcome of their choices. Task phases are described as based on definitions in Figure 2; correct choices are choices of the option with the higher probability of positive feedback, regardless of whether positive feedback was received; incorrect choices are choices with the higher probability of negative feedback. Positive feedback is the gain of 50 points irrespective of the participant’s choice, negative feedback is the loss of 50 points irrespective of the participant’s choice.

|  | Initial learning |  | Re-learning |  |  | Stability |  |
| --- | --- | --- | --- | --- | --- | --- | --- |
|  | Correct | Incorrect | Correct | Incorrect |  | Correct | Incorrect |
| Win | CI |  | CRe |  |  | CSt |  |
| Loss |  | II |  | Re | FRe | PESt | ISt |
|  |  |  |  | Re_par |  |  |  |

Table 1 Definitions of trial types within the task used for the analysis of functional magnetic resonance imaging data based on the task phase, the participant's choice, and the outcome of their choices.
| Abbreviation | Description |
| --- | --- |
| CI | Correct response and positive feedback during initial learning |
| CRe | Correct response and positive feedback during re-learning |
| CSt | Correct response and positive feedback during stability |
| CR = (CI + CRe + CSt) | All correct response and positive feedback trials |
| II | Incorrect response and negative feedback during initial learning |
| Re | Reversal error – Incorrect response and negative feedback during re-learning (excluding the last Re trial for each learning event) |
| Re_par | Reversal error – As above, but with EV the parametrically modulated such that activation decreases from 1 to (excluding) 0 in each learning event. |
| FRe | Final reversal error – The last incorrect response and negative feedback during re-learning (for each learning event) |
| PESt | Probabilistic error - Correct response but negative feedback during stability |
| ISt | Incorrect response and negative feedback during stability |

**Figure 2.**
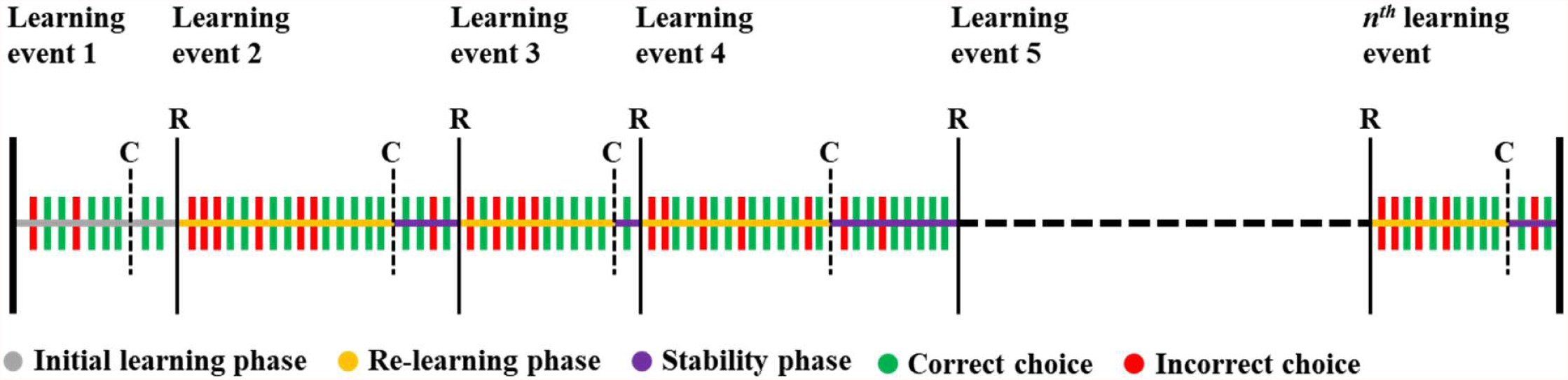
Trial and task phase overview of the serial reversal learning task. Dashed vertical lines show when criterion was reached (C); thin vertical lines show where outcome contingencies reversed (R) and a new learning event starts. Initial learning is the first learning event. After each reversal (R) participants are in the re-learning phase until they reach criterion (C). Participants are then in the stability phase until outcome contingencies reverse (R). The learning criterion must be maintained during the stability phase before reward contingencies reverse. Incorrect choices during the re-learning phase are defined as reversal errors, and the last reversal error of each re-learning phase is defined as the final reversal error. Each participant completes a total of 360 trials.

#### Model 2 – epoch-wide modelling across task phases for PPI analysis

The second general linear model modelled each epoch of the trial as a separate regressor and was used as the basis for our psychophysiological interaction analyses (PPI). These epochs were the decision-making phase, the anticipation phase, the outcome phase, and the cumulative total phase for each trial (see Figure 1 for a schematic of the trial epochs). The decision-making phase was subdivided into decision making during initial learning, re-learning and the stability phase. The outcome phase was subdivided into positive, negative, and neutral (i.e. no choice on that given trial) feedback during each of the initial learning, re-learning, and stability phases separately.

The purpose of the PPI analysis was to interrogate the functional connectivity of regions of interest in the orbitofrontal cortex and the thalamus with specific subregions of the striatum (Friston et al., 1997; O’Reilly et al., 2012).

Unilateral seed timeseries for the medial and lateral portions of the orbitofrontal cortex, and the mediodorsal and centromedian-parafascicular nuclei of the thalamus were extracted using subject-space masks generated using FreeSurfer, and subjects pre-processed BOLD data. Medial orbitofrontal cortex was included as a control region for the lateral orbitofrontal cortex due to their dissociable roles in reversal learning. The mediodorsal thalamus was included as a control region as although it has some projections to the striatum (which do not project to cholinergic interneurons), the cortex is its main target. Conversely, the centromedian-parafascicular nuclei project preferentially to the striatal cholinergic interneurons, and these projections have been implicated specifically in reversal learning (Bell, Langdon, et al., 2019). The centromedian-parafascicular nuclei and the mediodorsal nucleus also appear to have distinct functional roles during reversal learning. Previous evidence suggests the mediodorsal nucleus is important for using recent reward history to guide behaviour and minimising perseveration (Chakraborty et al., 2016), while the centromedian-parafascicular nuclei are important for generating multiple concurrent representations of contingencies that are relevant in different contexts, signalling behaviourally relevant sensory stimuli that signal a change in context, and minimising regressive error (Bradfield & Balleine, 2017; Brown et al., 2010; Matsumoto et al., 2001; Schepers et al., 2017; Yamanaka et al., 2018).

For each general linear model, the timeseries of a seed region was included as a regressor, and the interaction between the psychological regressors and the seed timeseries was calculated. Therefore, each general linear model contained sixteen regressors. Each permutation of psychological regressor and seed timeseries was run as a separate model; these are summarised in Table 2. This resulted in forty-eight PPI models for each subject. Contrast estimates were generated using the interaction term from each PPI to identify differences in functional connectivity for positive and negative feedback during the outcome epoch and differences in functional connectivity between the re-learning and stability phases for the decision making and feedback epoch. Region of interest analysis was used to restrict our PPI results to three functional subdivisions of the striatum (associative, limbic and motor) that were defined *a priori* as target areas (Bell, Langdon, et al., 2019; Choi et al., 2012). Regions of interest were ipsilateral to the seed region used for PPI analysis, based on the predominant anatomical connectivity of corticostriatal and thalamostriatal circuits (Bradfield, Bertran-Gonzalez, et al., 2013; Gourley et al., 2013).

**Table 2.**
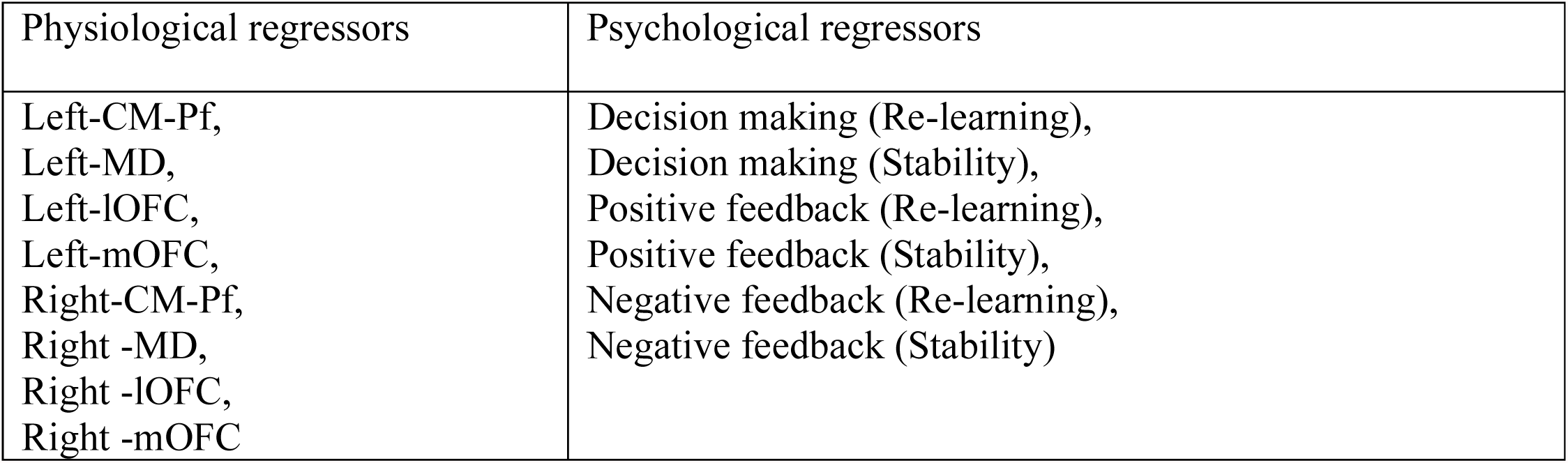
Overview of physiological and psychological regressors used in the general linear model for psychophysiological interaction analysis.

#### Group-level Estimates

Higher analysis was used to calculate group level activation for our contrast estimates. Age was included as a covariate of no interest in our higher level analysis to control for maturational effects on our results, given the age range of our participants (Boehme et al., 2017); the number of reversals was also included to control for the number of learning events each participant completed. Group estimates were calculated using FLAME 1 (FMRIB’s Local Analysis of Mixed Effects) in FSL using familywise error corrected (FWE) cluster thresholding (Z = 2.3, p < 0.05). For whole-brain contrasts, global and local maxima within each cluster were identified using a custom python script (https://github.com/bwilliams96/pyCL/blob/master/Label_copes.ipynb) that used atlasquery in FSL to produce an html report of the most probable region using the Harvard-Oxford cortical subcortical structural atlases. For each region, the coordinates, size, and p(FWE) corresponding to the highest z-value are reported. Our results follow published reporting guidelines for functional neuroimaging studies (Poldrack et al., 2008).

## Results

### Behavioural summary

All subjects included in the analysis experienced an average of 24.62 (SD = 5.67; Range = 10 - 37) reversals during the task and selected an image on >95% of trials. Correct responses, regardless of outcome, were made at significantly greater than chance level (mean correct choices = 254.38, 95% CI [248.66, *∞], t*(28) = 22.13, *p* < .001, SD = 18.10, Range = 193 - 290), and participants experienced an average of 24.62 (SD = 5.77; Range = 10 - 37) reversals (Figure 3). The average number of trials taken to reach criterion was 8.76 (SD = 4.33; Range = 5 - 40); an average of 3.07 (SD = 2.30; Range = 0 - 18) perseverative errors were made following the reversal of contingencies before reaching criterion in each learning event. Participants did not respond on an average 2.79 trials (SD = 3.59; Range = 0 - 15) during the task. An average of 4,718.97 (SD = 1,065.31, Range = 1200 - 6800) points were collected by the end of the task. The average reaction time of participants to choose following the onset of the stimuli was 621.57 milliseconds (SD = 123.86, Range = 367.07 - 917.30).

**Figure 3.**
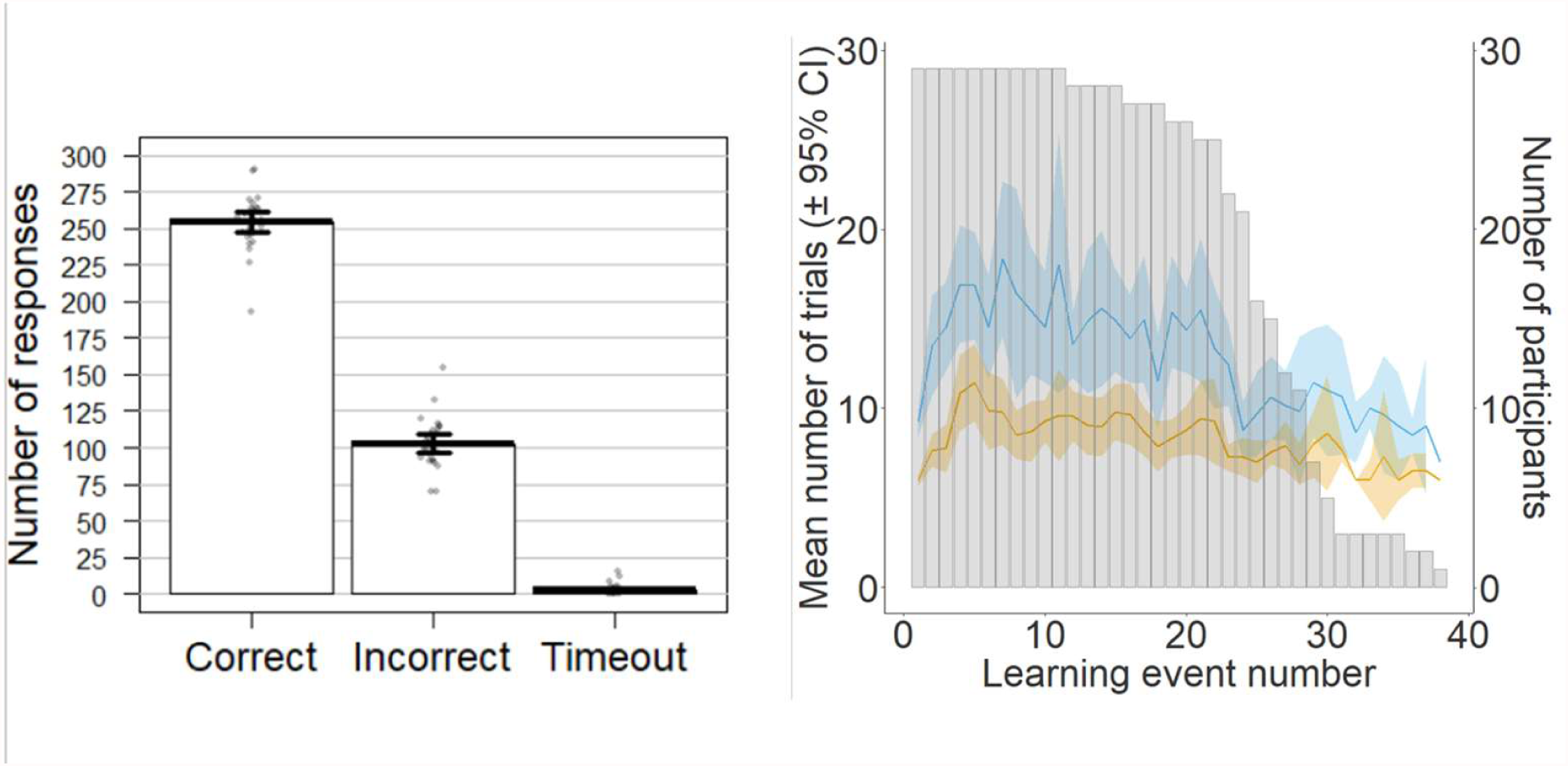
Mean (±95% confidence intervals) number of correct, incorrect, and missed responses for each participant across the task. Mean (±95% confidence intervals) number of trials (blue) and trials to criterion (orange) in each learning event; grey bars are the number of participants who reached each learning event.

### fMRI results

#### Task-related activations are in line with previous studies of probabilistic reversal learning

First, we wanted to understand whether our data was in line with previous reports of probabilistic reversal learning. If results from these analyses are aligned with previous reports, then this suggests our task is broadly comparable with other designs. Significant bilateral insula activation was found when contrasting final reversal errors and correct responses (FRe > CR), in line with previous findings (Table 3, Figure 4) (Cools et al., 2002; Dodds et al., 2008; Freyer et al., 2009; Waegeman et al., 2014; Yaple & Yu, 2019; Zeuner et al., 2016). Reversal errors that did not lead to a change in behaviour (Re > CR) showed activation consistent with past findings in the anterior cingulate/paracingulate cortex (Kringelbach & Rolls, 2003), bilateral insular cortex and right frontal operculum cortex/inferior frontal gyrus (Table 3) (Mitchell et al., 2008).

**Table 3.**
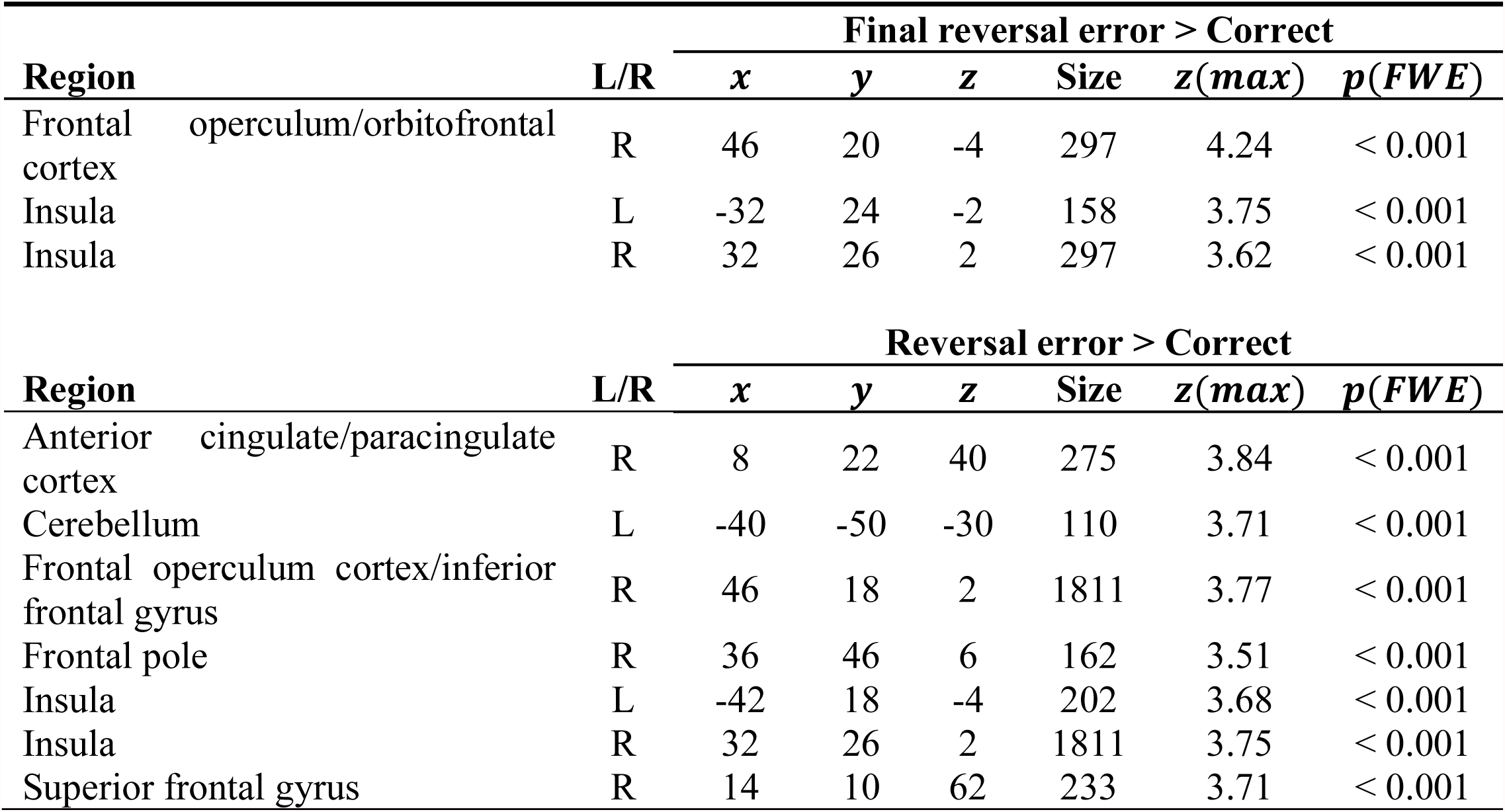
Significant clusters from whole brain analysis of contrasts used in previous studies of probabilistic reversal learning (see text for details and citations).

**Figure 4.**
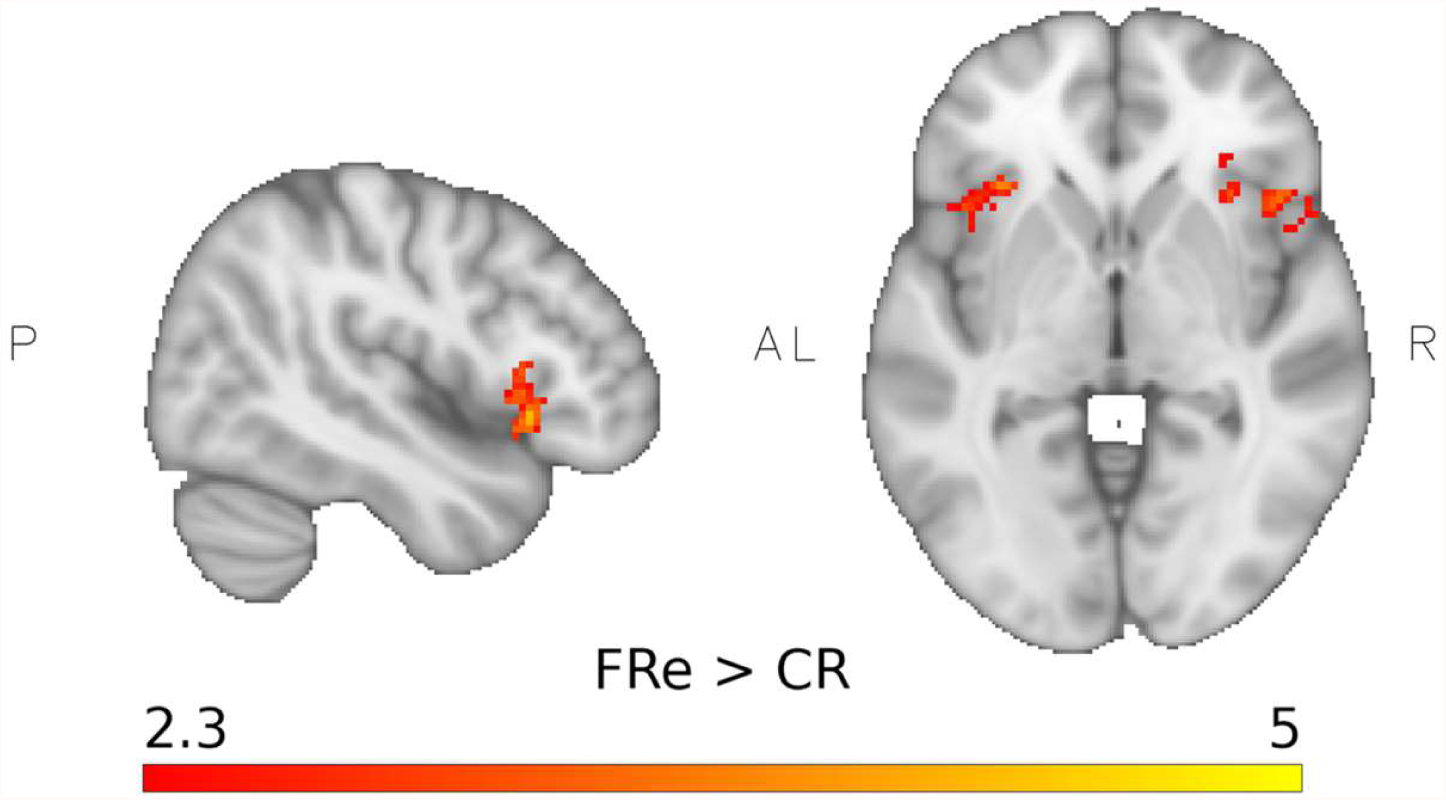
Whole brain analysis shows similar activations for final reversal errors versus correct response as in previous studies of reversal learning (see text for details). Significant clusters were identified in bilateral insula (peak coordinates were *x* = −32, *y* = 24, *z* = −2, *z*(*max*) = 3.75 for the left insula; *x* = 32, *y* = 26, *z* = 2, *z*(*max*) = 3.75 for the right insula) and in the right orbitofrontal cortex (peak coordinates *x* = 46, *y* = 20, *z* = −4, *z*(*max*) = 4.24). A significance threshold of *p*(*FWE*) < 0.05 and a cluster threshold of z > 2.3 was used.

#### Centromedian-parafascicular nuclei and lateral orbitofrontal cortex show increased functional connectivity with striatal regions during the processing of negative feedback

We were next interested in assessing functional connectivity between the orbitofrontal cortex and striatum, and between the thalamus and the striatum. To do this we use psychophysiological interaction analysis. We used unilateral seed regions in the medial and lateral orbitofrontal cortex, and in the centromedian-parafascicular and mediodorsal nuclei of the thalamus. We measured ipsilateral functional connectivity with the striatum and restricted our analysis to regions of the associative dorsal striatum, motor dorsal striatum, and limbic ventral striatum (Bell, Langdon, et al., 2019; Choi et al., 2012). For the outcome epoch we assessed differences in functional connectivity between the re-learning and stability phases, and between positive and negative feedback. For the decision-making epoch we assessed differences in functional connectivity between the re-learning and stability phases (Table 2).

##### Outcome valence

Functional connectivity between the left centromedian-parafascicular nucleus of the thalamus and the associative dorsal striatum during the outcome epoch was significantly greater for negative feedback than positive feedback (z(max) = 3.57, MNI coordinates = [-26, 4, -2], 38 voxels, p = 0.012, Figure 5B & C). We also found significantly greater functional connectivity for negative feedback than positive feedback between the right lateral orbitofrontal cortex, and the associative dorsal striatum (z(max) = 4.14, MNI coordinates = [18, 16, -4], 39 voxels, p = 0.008, Figure 5E). Increased functional connectivity for negative feedback, relative to positive feedback suggests that thalamostriatal and corticostriatal circuits may use negative outcomes to guide adaptive behaviour during serial reversal learning.

**Figure 5.**
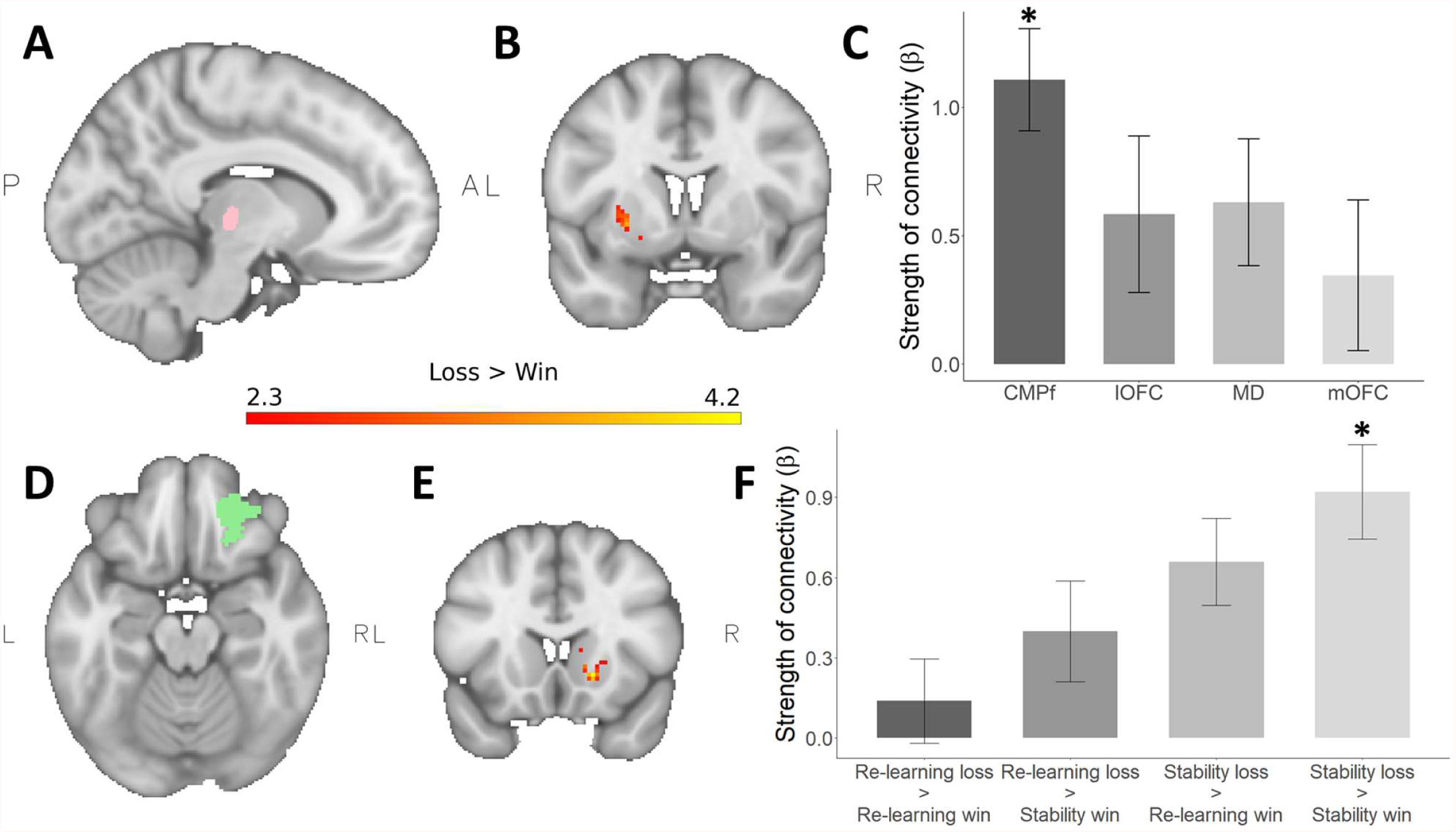
A: Left centromedian-parafascicular (CMPf) seed region. B: Functional connectivity between the left centromedian-parafascicular and the left associative striatum was significantly greater during the processing of negative feedback versus positive feedback (peak coordinates *x* = −26, *y* = 4, *z* = −2, *z*(*max*) = 3.57). No significant differences in this contrast were seen when comparing between phases of the task. C: Strength of functional connectivity between left cortical (lOFC: lateral orbitofrontal cortex, mOFC: medial orbitofrontal cortex) and thalamic (CMPf, MD: mediodorsal nucleus) seeds with the associative dorsal striatum for negative feedback versus positive feedback. Significant functional connectivity was observed between the left centromedian-parafascicular nuclei and the striatum, but other regions did not show significant functional connectivity. D: Right lOFC seed region. E: Functional connectivity between the right lOFC and the right associative striatum was significantly greater during the processing of negative feedback versus positive feedback (peak coordinates *x* = −18, *y* = 16, *z* = −4, *z*(*max*) = 4.14). F: Significant differences in functional connectivity were observed for negative versus positive feedback during the stability phase, but not between other phases of the task. A significance threshold of *p*(*FWE*) < 0.05 and a cluster threshold of z >2.3 was used.

To determine whether differences in functional connectivity between the centromedian-parafascicular nucleus or the lateral orbitofrontal cortex and the associative striatum were localised to a specific phase of the task we calculated differences in functional connectivity for negative and positive feedback across the re-learning and stability phases of the task. Functional connectivity between the lateral orbitofrontal cortex and the associative striatum was significantly greater for negative feedback than positive feedback during the stability phase (z(max) = 3.21, MNI coordinates = [14, 16, 0], 40 voxels, p = 0.006, Figure 5F). No differences were found between the centromedian-parafascicular and the associative striatum for negative and positive feedback across re-learning and stability. These findings are suggestive of a general error signal between the centromedian-parafascicular and the associative striatum used to guide behaviour by signalling potential changes in context based on negative feedback. Conversely, error signals between the lateral orbitofrontal cortex and associative striatum may be specifically used to implement a change in response strategy, in line with previous literature (Hampshire et al., 2012; Rygula et al., 2010). Importantly, no differences in functional connectivity between the medial orbitofrontal cortex and mediodorsal thalamus with our striatal regions of interest was observed during the outcome epoch of the task. These regions were included as control regions, and support the specificity of the thalamus with the dorsal striatal cholinergic system, and the lateral orbitofrontal cortex during serial reversal learning. The dorsal striatal cholinergic system is preferentially innervated by the centromedian and parafascicular nuclei, while the mediodorsal nucleus has few projections to the striatal cholinergic system (Smith et al., 2009). The striatal cholinergic system has an important role during reversal learning, and is associated with the generation and flexible use of multiple internal representations (Bell et al., 2018; Bradfield, Bertran-Gonzalez, et al., 2013; Bradfield & Balleine, 2017; Brown et al., 2010; Ragozzino et al., 2009; Stalnaker et al., 2016). Therefore, although we have not measured striatal cholinergic activity directly, the specificity of these results suggests that functional connectivity between the centromedian-parafascicular nucleus and the dorsal striatum might be associated with striatal cholinergic interneuron activity. Additionally, while the medial orbitofrontal cortex is involved in outcome evaluation and goal-directed behaviour, its inactivation is associated with general impairments in probabilistic learning (Dalton et al., 2016). Conversely, lateral orbitofrontal inactivation preferentially impairs reversal learning (Dalton et al., 2016), in line with other work suggesting the lateral orbitofrontal cortex is important for initiating change and minimising perseveration after the reversal of outcome contingencies (Bell, Langdon, et al., 2019; Hampshire et al., 2012; Hervig et al., 2020). Therefore, the specificity of lateral orbitofrontal result suggests its functional connectivity with the dorsal striatum changed as a function of reversal learning, while no differences in functional connectivity for the medial orbitofrontal cortex suggests the strength of its connectivity was consistent across the task.

##### Decision-making

Next, we assessed functional connectivity between our seed regions in the orbitofrontal cortex and thalamus with the striatum during decision making. Functional connectivity between the right mediodorsal thalamus and the ventral striatum was significantly greater during the decision-making epoch of the re-learning phase than the stability phase (z(max) = 3.03, MNI coordinates = [16, 6, - 12], 8 voxels, p = 0.046).

Although the mediodorsal thalamus was included as control regions based on the specificity of centromedian-parafascicular connectivity with the dorsal-striatal cholinergic system, there is prior evidence from animal and human literature for a role for the mediodorsal thalamus and ventral striatum in reversal learning. For instance the ventral striatopallidal circuit, which includes the ventral striatum and mediodorsal thalamus, helps the inhibition of responding to previously rewarded stimuli after contingency reversal, as lesions to the circuit impair reversal learning, but not stimulus discrimination or simple stimulus-outcome association learning (Ferry et al., 2000; Price, 2005).

In the present study, increased coherence between the mediosorsal thalamus and ventral striatum during decision making may help prevent perseverative errors during re-learning, i.e. after the reversal has been instigated. We ran supplementary analyses to test this hypothesis but found no correlation between participant’s average number of perseverative errors and the difference in functional connectivity between the re-learning and stability phases (*r* = -0.029, *p* = 0.881).

No differences in functional connectivity during decision making between the orbitofrontal cortex or centromedian-parafascicular nucleus and the striatum were observed during re-learning and stability phases.

#### Cortical error signals are parametrically modulated by response perseveration

Following our findings describing the role of thalamostriatal and corticostriatal connectivity in reversal learning, we next ran a series of whole brain exploratory analyses. These analyses aimed to investigate further how negative and positive feedback may guide adaptive behaviour during serial reversal learning; we modelled different trial types as individual explanatory variables (Table 1).

We first tested for error signals that could relate to a change in behavioural strategy. To do this we compared final reversal errors, as defined by Cools et al. (2002), with errors unrelated to reversal learning. Specifically, we compared final reversal errors with incorrect choices during initial learning (FRe > II) and found significant increases in activation in the orbitofrontal and insular cortices and middle frontal, paracingulate, and superior frontal gyri (Table 4). However, these differences do not provide causal evidence of activation resulting in behavioural change, since the difference could merely be due to differences in responding to negative outcomes during initial and reversal learning. Therefore, we compared error signals leading to a change in strategy with those that did not (FRe > II) > (Re > II) to test whether these effects were specifically associated with strategy change. None of our initial clusters survived thresholding. One potential explanation for this lack of overall statistical effect is that evidence leading to a change in strategy is accumulated gradually. Therefore, this gradient could be averaged out when grouping trial types.

**Table 4.** Significant clusters from whole brain analysis of contrasts that describe how perseverative error processing may be used to guide adaptive behaviour following the reversal of outcome contingencies.

| Region | L/R | Final reversal error $>$ Incorrect Initial learning | | | | | |
| --- | --- | --- | --- | --- | --- | --- | --- |
| | | $x$ | $y$ | $z$ | Size | $z(max)$ | $p(FWE)$ |
| Angular gyrus | R | 52 | -46 | 58 | 477 | 3.79 | $< 0.001$ |
| Frontal operculum cortex | L | -40 | 18 | 0 | 163 | 3.68 | 0.001 |
| Frontal operculum/orbitofrontal cortex | R | 44 | 20 | -4 | 497 | 4.71 | $< 0.001$ |
| Inferior frontal gyrus | R | 52 | 20 | -4 | 497 | 3.78 | $< 0.001$ |
| Insula | L | -34 | 20 | -2 | 163 | 3.36 | 0.001 |
| Insula | R | 32 | 26 | 4 | 497 | 3.85 | $< 0.001$ |
| Lateral occipital cortex | R | 42 | -58 | 56 | 477 | 3.74 | $< 0.001$ |
| Middle frontal gyrus | R | 46 | 24 | 44 | 122 | 3.23 | 0.013 |
| Orbitofrontal cortex | L | -32 | 24 | -4 | 163 | 3.86 | 0.001 |
| Paracingulate gyrus | R | 2 | 14 | 52 | 464 | 4.32 | $< 0.001$ |
| Superior frontal gyrus | R | 2 | 28 | 52 | 464 | 3.91 | $< 0.001$ |
| Supramarginal gyrus | R | 56 | -44 | 50 | 477 | 4.12 | $< 0.001$ |

| Region | L/R | Reversal error Parametric modulation $>$ Incorrect Initial learning | | | | | |
| --- | --- | --- | --- | --- | --- | --- | --- |
| | | $x$ | $y$ | $z$ | Size | $z(max)$ | $p(FWE)$ |
| Central opercular cortex | L | -58 | -6 | 8 | 162 | 3.43 | $< 0.001$ |
| Central opercular cortex | R | 56 | 2 | 4 | 375 | 3.70 | $< 0.001$ |
| Heschl's gyrus | L | -46 | -24 | 12 | 120 | 3.47 | 0.003 |
| Heschl's gyrus | R | 44 | -26 | 12 | 416 | 3.63 | < 0.001 |
| Insula | R | 40 | -16 | 6 | 375 | 3.50 | < 0.001 |
| Lateral occipital cortex | L | -52 | -70 | 36 | 92 | 2.77 | 0.024 |
| Lingual gyrus | L | -30 | -56 | 4 | 192 | 4.56 | < 0.001 |
| Paracingulate gyrus | L | 0 | 44 | -8 | 327 | 4.38 | < 0.001 |
| Parietal operculum cortex | L | -52 | -26 | 12 | 120 | 2.77 | 0.003 |
| Parietal operculum cortex | R | 56 | -24 | 18 | 416 | 3.86 | < 0.001 |
| Planum temporale | L | -64 | -24 | 14 | 184 | 2.96 | < 0.001 |
| Postcentral gyrus | R | 66 | -12 | 14 | 416 | 3.58 | < 0.001 |
| Posterior cingulate cortex | L | -4 | -52 | 14 | 191 | 3.71 | < 0.001 |
| Precentral gyrus | L | -58 | 6 | 2 | 162 | 3.43 | < 0.001 |
| Precentral gyrus | R | 62 | 4 | 6 | 375 | 3.49 | < 0.001 |
| Precuneous | L | -8 | -56 | 26 | 191 | 3.81 | < 0.001 |
| Subcallosal cortex | L | 0 | 22 | -14 | 327 | 3.47 | < 0.001 |
| Supramarginal gyrus | L | -66 | -24 | 26 | 184 | 3.34 | < 0.001 |
| Supramarginal gyrus | R | 62 | -28 | 24 | 416 | 4.02 | < 0.001 |

To test this hypothesis, we parametrically modulated reversal errors not leading to a change in strategy (Re_par), scaled descending from one to (excluding) zero. Re_par was then de-meaned, allowing us to combine learning events of differing lengths. We used a descending parametric modulator because we expect early reversal errors to produce greater responsivity than late reversal errors, akin to a prediction error signal. Reversal errors showed significant parametric modulation in regions involved in error processing, including the posterior insula, Heschl’s gyrus, and the posterior cingulate cortex (Table 4). These findings suggest activation may be related to the accumulation of evidence to determine when to reverse.

### Prefrontal regions supporting the implementation of cognitive flexibility

Considering our findings describing how errors may be used to guide behaviour, we next wanted to explore differences in activation for choices made during the re-learning and stability phases. We contrasted trials where participants made correct responses during the stability versus the re-learning phase (CSt > CRe) and found a single significant cluster in the left frontal pole (Table 5, Figure 6). During the stability phase, participants demonstrate behaviourally that they understand which is the choice that is most likely to lead to a positive outcome. However, the relationship between actions and outcomes may be more uncertainty during the re-learning phase, given the recent reversal of reward contingencies. It has been suggested that the frontal polar cortex tracks the relative advantage of alternative response strategies, and recruits prefrontal regions to shift behaviour when the alternative strategy becomes advantageous (Mansouri et al., 2017). Therefore, while participants use a single response strategy during stability to gain positive outcomes, they may use the frontal polar cortex to track the relative advantage of the unchosen option.

**Table 5.**
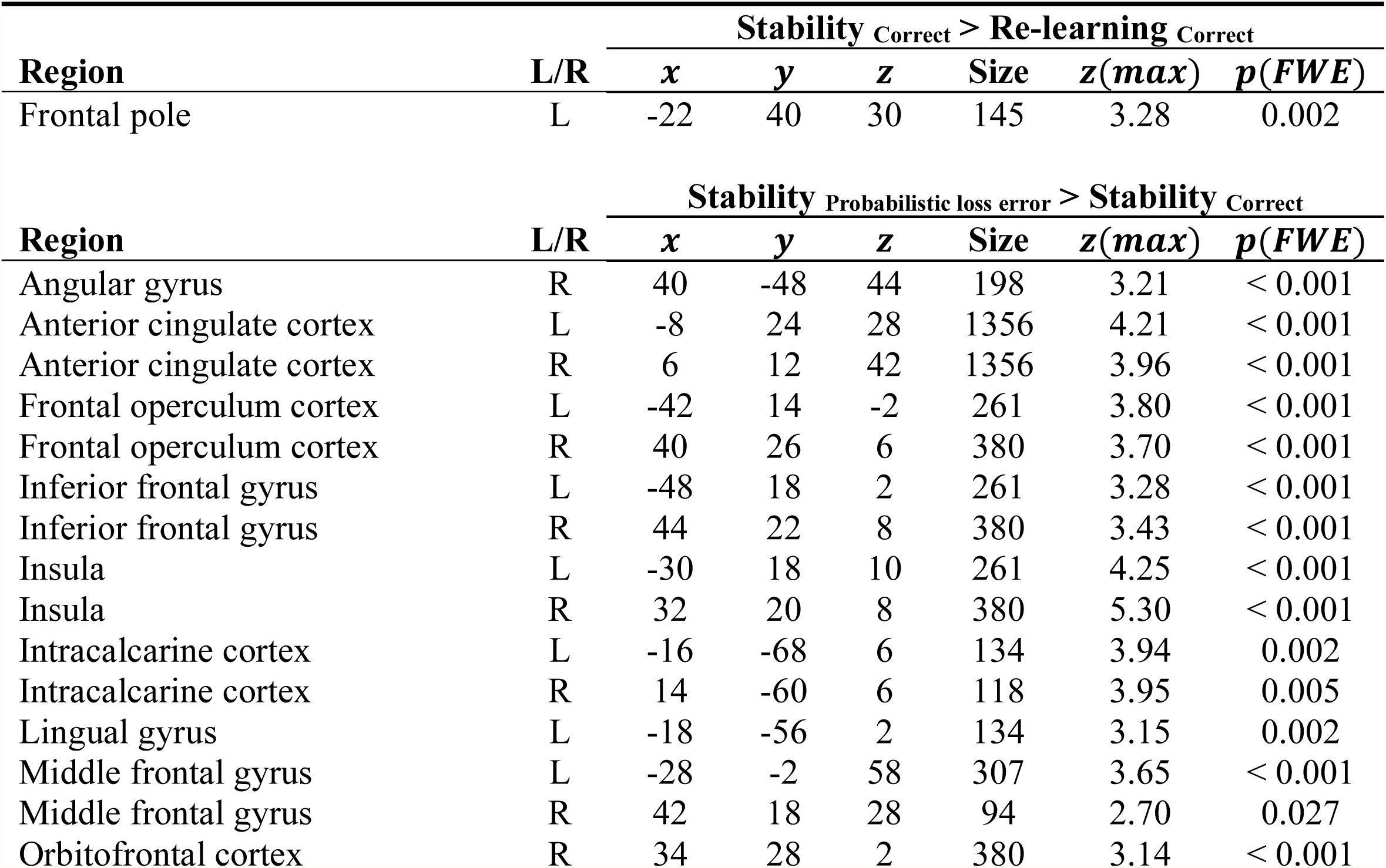
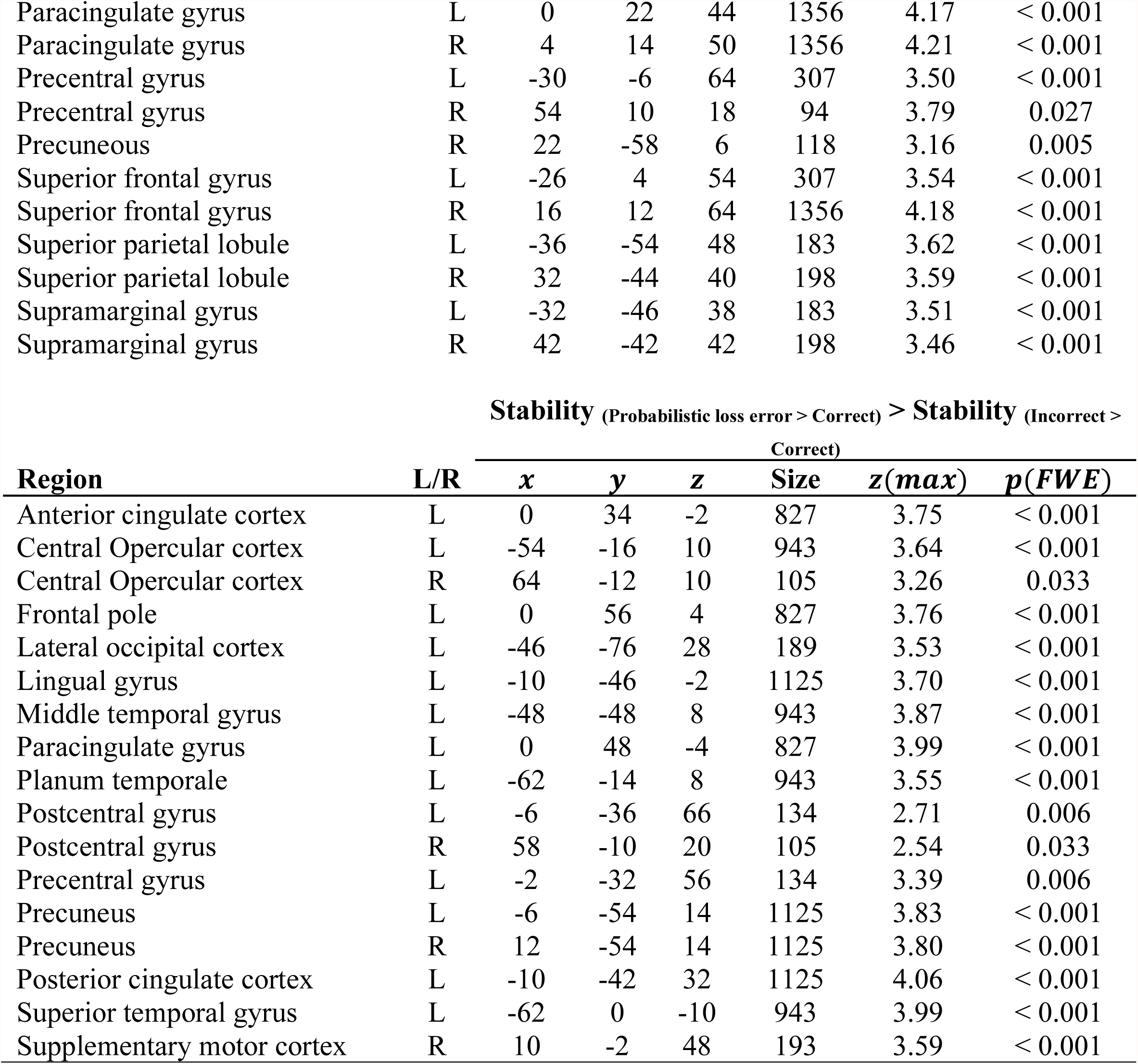
Significant clusters from whole brain analysis of contrasts showing differences between making correct choices at during different phases of the task, and how unexpected feedback may be used to prepare switching behaviour in response to negative feedback.

**Figure 6.**
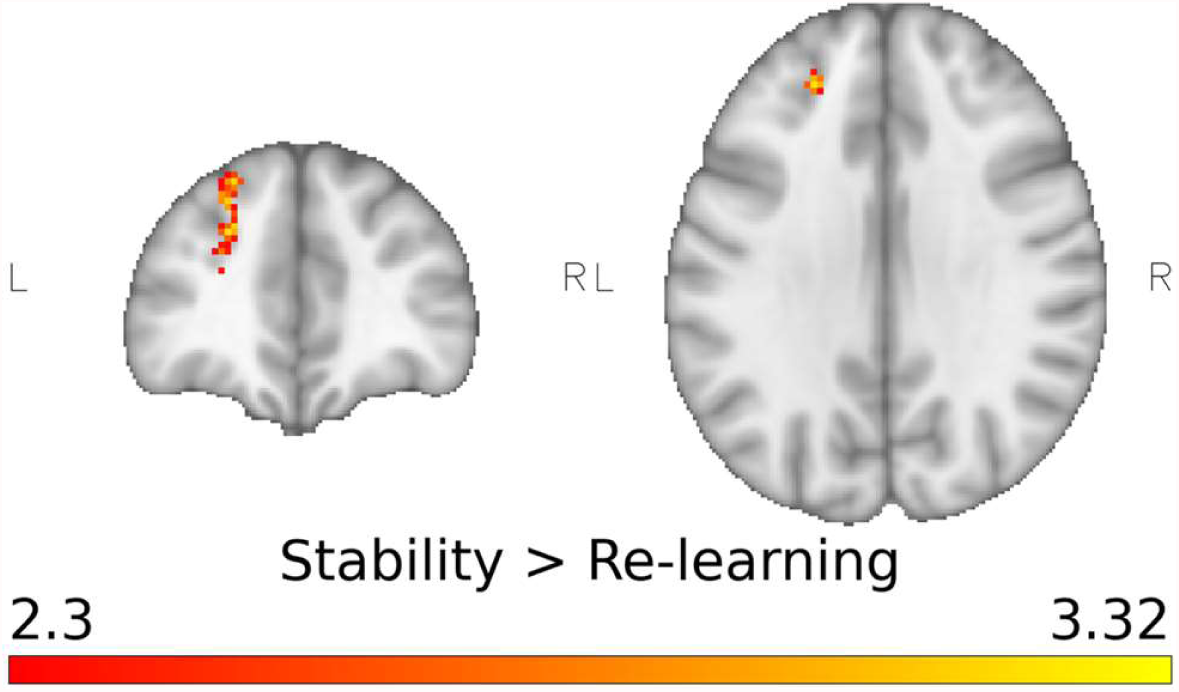
The frontal pole showed significantly increased activation when participants made correct choices during the stability phase versus re-learning (peak coordinates *x* = −22, *y* = 40, *z* = 30, *z*(*max*) = 3.28). This signal may reflect tracking the reliability of the alternative task strategy during reversal learning. Participants have demonstrated behaviourally that they understand the currently correct action policy once they have reached the stability phase, whereas the correct policy would be more ambiguous during re-learning, and therefore there is not a clear correct and alternative action policy. A significance threshold of *p*(*FWE*) < 0.05 and a cluster threshold of z >2.3 was used.

Outcomes that deviate from expectations are also likely to guide behaviour given that correct and incorrect choice are mutually exclusive in our task. Therefore, we next compared activation when feedback was congruent or incongruent with a participant’s expectations by contrasting correct choices during the stability phase that led to negative versus positive feedback (PESt > CSt). This type of negative feedback, i.e. following a choice they considered correct, could indicate to the participant that reward contingencies changed as they are incongruent with expectations. The contrast showed increased activation across the cortex, including regions involved in feedback monitoring such as the angular gyrus, anterior cingulate, inferior frontal gyrus, insula, and orbitofrontal cortex (Table 5, Figure 7A). To confirm the specificity of this result we looked for the effect over and above activation differences due to positive and negative feedback (PESt > CSt) > (ISt > CSt). Activation in several regions discriminated between incongruent negative feedback and congruent positive feedback for correct responses during the stability phase, over and above responsivity to punishment (Table 5, Figure 7B). Most notably, significant activation was consistently found in the anterior cingulate and paracingulate cortex for activation over and above differences due to positive and negative feedback; significant activation in the lingual and precentral gyri was also observed in both analyses ((PESt > CSt) > (ISt > CSt) and PESt > CSt). Anterior cingulate has been found to support adaptive behaviour in macaques by using errors to guide choice, while lesions to the anterior cingulate led to impairments in reversal learning (Chudasama et al., 2013).

**Figure 7.**
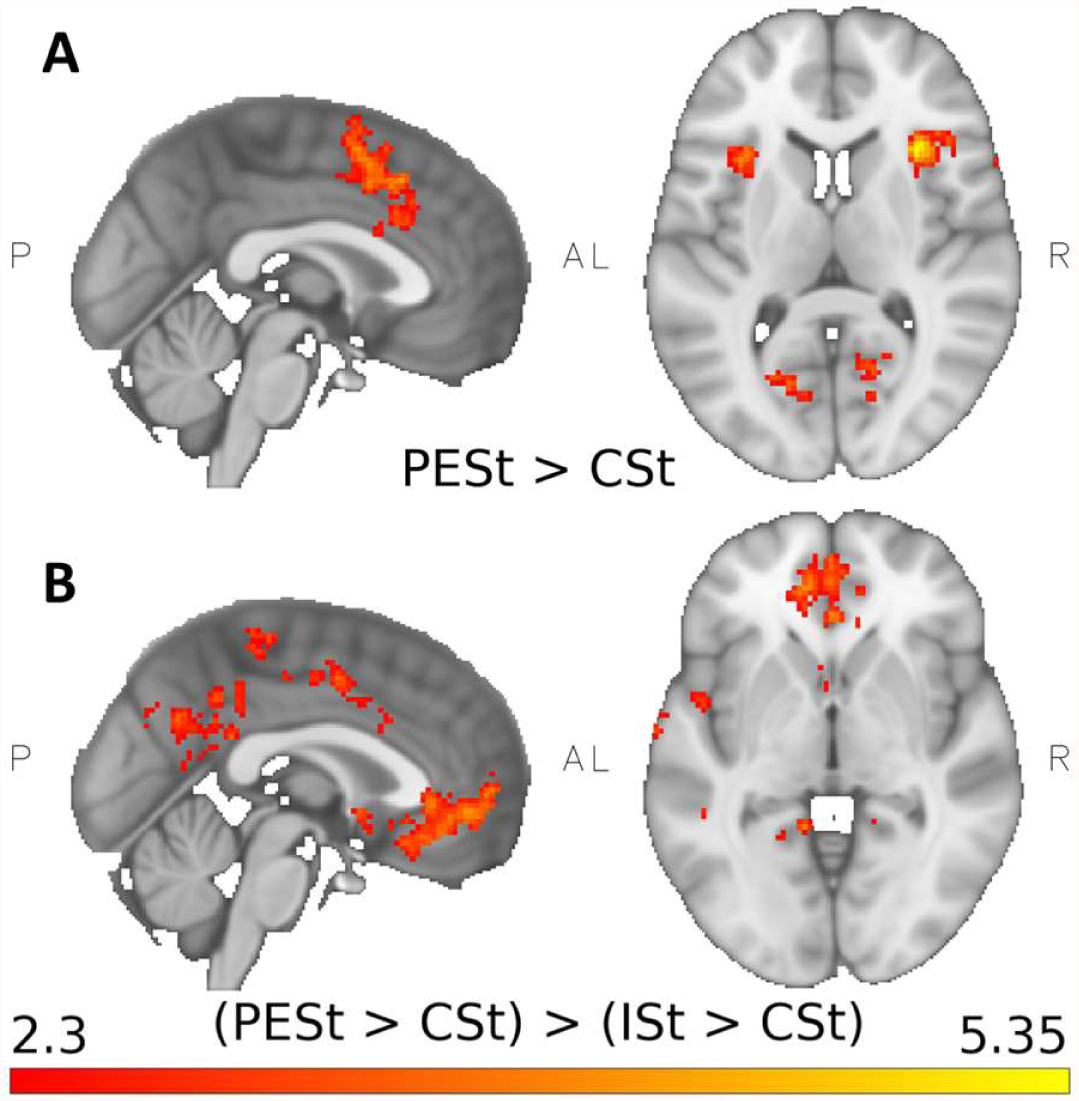
A: Significantly greater activation for probabilistic loss errors than correct choices during the stability phase was seen in the anterior cingulate, orbitofrontal cortex, insula, and inferior frontal gyrus (full details on regions and cluster statistics in Table 5). This contrast shows differences in activation when actual outcomes are incongruent with expected outcomes and may indicate regions that are involved in signalling unexpected outcomes that could lead to a change in behaviour. B: Significant differences in activation for probabilistic loss errors than correct choices, above activation for negative versus positive feedback (Probabilistic loss error > correct) > (Incorrect > correct) (all stability). Several regions, including the anterior cingulate cortex (see Table 5 for full details) show increased activation to probabilistic loss errors specifically. These regions may use probabilistic loss errors to signal a potential change in context that could lead to the reversal of behaviour. A significance threshold of *p*(*FWE*) < 0.05 and a cluster threshold of z >2.3 was used.

## Discussion

The present study aimed to investigate the influences of cortical and thalamic brain regions on striatal activity during probabilistic reversal learning. We found that functional connectivity between the centromedian-parafascicular nuclei of the thalamus and a portion of the associative striatum was significantly greater during the processing of negative compared to positive feedback. A similar pattern of functional connectivity was observed between the lateral orbitofrontal cortex and the associative striatum. These results suggest the centromedian-parafascicular nuclei, and the dorsal striatum use negative feedback as a general error signal to promote flexibility. Lateral orbitofrontal cortex and dorsal striatum connectivity was specific to when participants were using the alternative response strategy, suggesting connectivity may be involved in the implementation of change to an alternate strategy. Additionally, we explored how cortical activity may influence cognitive flexibility. Following the reversal of contingencies, parametric modulation of regions involved in error processing, including the posterior cingulate and insula cortices, and Heschl’s gyrus, was observed. This modulation may be related to the gradual accumulation of evidence that contingencies have reversed from negative feedback in preparation for change. Orbitofrontal and insular cortices and middle frontal, paracingulate, and superior frontal gyri showed significant activation for probabilistic loss errors during the stability phase. These regions have previously been reported to be important for error processing and change detection, and therefore activation may be related to anticipating future reversals. Lastly, we provide evidence suggesting that the frontal polar cortex is involved in monitoring the relative advantage of alternate response strategy, and a set of regions including the inferior frontal gyrus, anterior cingulate and orbitofrontal cortices that may signal when a reversal has occurred.

Connectivity between the centromedian-parafascicular nuclei and striatum has previously reported to be involved in the expression of adaptive behaviour in rodents and humans (Bell, Langdon, et al., 2019; Bradfield, Bertran-Gonzalez, et al., 2013; Bradfield, Hart, et al., 2013; Brown et al., 2010; Yamanaka et al., 2018). Here we provide further evidence for the premise that thalamostriatal circuits are important for adaptive behaviour by showing that functional connectivity is significantly greater during the processing of negative versus positive feedback in a task where behavioural adaptation relies on the reliable tracking of negative outcomes. This difference in functional connectivity is in line with the purported role of the centromedian-parafascicular in signalling contextual change (Bradfield, Bertran-Gonzalez, et al., 2013; Yamanaka et al., 2018). In a simple, two-choice task, actual outcomes are likely to match expected outcomes; therefore, incongruent feedback might suggest that outcome contingencies had reversed. Thus, in this task, the centromedian-parafascicular may be involved in detecting changes in behavioural context by tracking negative feedback. This information could then be used to infer that the current behavioural policy needs to be changed.

Indeed, this suggestion is in line with previous research showing greater activation in the centromedian-parafascicular nuclei when overcoming bias and responding to unexpected outcomes (Matsumoto et al., 2001; Minamimoto et al., 2005, 2014). Centromedian-parafascicular activity may indicate a general error signal that is used to guide behaviour by signalling a potential change in context following negative outcomes. This may also explain why significant differences in connectivity were not found during feedback in different phases of the task, i.e. thalamostriatal connectivity did not discriminate between losses and wins during stability or re-learning. This is in line with previous work showing that centromedian-parafascicular neurons habituate to non-reward but not reward-related stimulation (Alloway et al., 2014; Matsumoto et al., 2001). If we did see differences between phases of the task, then this would suggest the centromedian-parafascicular nuclei, and the dorsal striatum have reduced connectivity during some phases compared to others.

Conversely, the lateral orbitofrontal cortex and the striatum show increased connectivity during negative feedback, but only during the stability phase, suggesting this connectivity may support changes in behaviour following reversal. This finding is in line with previous evidence that suggests the lateral orbitofrontal cortex is involved in the implementation of changes in behaviour during reversal learning (Hampshire et al., 2012; Rygula et al., 2010).

Thalamic and cortical connections may be integrated within the striatum to coordinate cognitive flexibility. For instance, thalamostriatal connections have the potential to signal changes in context to the cholinergic interneurons at any time, simply by virtue of a non-specific error signal. Meanwhile, corticostriatal connections may either attenuate or enhance the influence of this thalamostriatal input to the striatum. For instance, other external inputs to the orbitofrontal cortex may signal that a change in behaviour is required, at which point connections from the orbitofrontal cortex to the striatum will modulate the influence of cholinergic interneurons on the output of the striatum.

Previous studies using reversal learning have indicated that quantitative activation differences exist between final reversal errors and errors not leading to a change in strategy (Cools et al., 2002; Culbreth et al., 2016; Remijnse et al., 2006), although, we did not find such differences in our data. One potential explanation is the differences in the design of our probabilistic learning task. For instance, Cools et al. (2002) had participants complete thirty minutes of probabilistic discrimination learning training, Culbreth et al. (2016) also had participants complete practice trials (though they do not describe the extent of practice), and instructed them to stick with a response. Therefore, participants in these studies received more training than our participants did. In terms of prior experience, the participants of Remijnse et al. (2006) are closest to our own, as they completed thirty trials of probabilistic discrimination learning without reversal. Yet, although Remijnse et al. (2006) used probabilistic feedback for correct choices, incorrect choices were always deterministic, and participants were told what stimulus would initially be correct. Therefore, discrimination of correct and incorrect choices should require less effort from the participant. By contrast, participants in our task were relatively naïve to probabilistic discrimination learning, and though they received instruction about probabilistic feedback and the existence of reversals they were not explicated instructed how to make choices in the task. We therefore cannot assume that reversal errors and final reversal errors in our task are necessarily equivalent with those of Cools et al. (2002), Culbreth et al. (2016) and Remijnse et al. (2006). Importantly, we also observe parametric modulation of reversal error signals in regions involved in change detection such as the insula and Heschl’s gyrus, and the posterior cingulate cortex. Furthermore, these differences in activation exist over and above negative feedback during initial learning. This may explain why Cools et al. (2002) did not see any modulatory effect of the number of preceding reversal errors on the final reversal error; here, we find that the insula appears to be modulated by preceding perseverative errors. Alternatively, the relatively small sample size of thirteen subjects used by Cools et al. (2002) could mean parametric modulation could not be detected in their dataset due to insufficient statistical power.

Model-based approaches to learning assume mental representations of an environment are used to guide goal-directed behaviour. This representation could include multiple plausible environmental contexts, with representation of the current context being driven by recent experience. The orbitofrontal cortex is associated with the representation of the current context (Schuck et al., 2016; Wilson et al., 2014); in particular medial orbitofrontal and ventromedial prefrontal activity is thought to be related to the reliability of the inferred context (Domenech & Koechlin, 2015). However, an environment with multiple contexts requires arbitration between them for an individual to respond adaptively to change. The frontal polar cortex, found exclusively in primates, is well placed to support such adaptive responding (Bunge, 2004; Bunge et al., 2005; Koechlin, 2014; Sakai & Passingham, 2006), and is associated with monitoring alternative strategies (Boorman et al., 2009). Here, we find increased activation in the frontal polar cortex during correct choices in stability compared to re-learning, and postulate that this is related to monitoring the reliability of the alternative task context. Though this study was not specifically designed to test this hypothesis, our reversal learning task is structured in such a way that once participants reach the stability phase, they have demonstrated behaviourally that they understand the current context of the task due to our stringent learning criterion. During the re-learning phase, the reversal of contingencies increases the relative uncertainty around the currently correct choice, making the likelihood for each choice more similar. Participants need to determine whether contingencies have reversed or not. Because either choice could currently be correct, there is no alternative context during re-learning. However, after reaching the stability phase participants have demonstrated behaviourally that they understand the current task context. Therefore, the frontopolar cortex can track the relative reliability of the alternative context, while the medial orbitofrontal cortex implements the behavioural policy associated with the current context during the stability phase. This proposition is supported by the error signals we found when participants received probabilistically incorrect feedback to correct choices during stability as compared to rewarded correct choices. Here we see increased activation in regions often associated with error feedback, such as the insula and inferior frontal gyrus, but we also see increased activation in the orbitofrontal and anterior cingulate cortices. Error-dependent changes in activity within the orbitofrontal cortex may indicate a decrease in the reliability of the current context, due to choice outcomes being incongruent with the estimate of the current context (Ghahremani et al., 2010). Alternatively, orbitofrontal cortex activity may indicate preparation to switch responding as is suggested by our functional connectivity results. Anterior cingulate activation may signal outcomes that are unexpected based on the current context and this salient event may increase attentional resources for monitoring outcomes with a view to potentially change strategy (Behrens et al., 2007; Chudasama et al., 2013; Liu et al., 2015).

There are several limitations with the current study which also provide possible avenues for future research. The first is the spatial specificity of our functional signal within subregions of the thalamus. In this study we aimed to minimise cross-contamination of our functional signal between nuclei within the thalamus by reducing our voxel size and smoothing kernel during preprocessing. Nevertheless, it is likely that signal blurring would still occur at the anatomical boundaries of nuclei within the thalamus, meaning that our timeseries used for psychophysiological interaction analysis may be influenced by more than one anatomical region. Therefore, it would be useful to validate these findings using ultra-high field magnetic resonance imaging as this would allow for greater spatial specificity and would provide further evidence that this signal is localised within the centromedian-parafascicular nuclei. Secondly, despite efforts taken to optimise echo-planar image acquisition to reduce dropout in the orbitofrontal cortex (Volz et al., 2019; Weiskopf et al., 2007), we found that for some participants there was still partial signal loss in the most rostral portions of the orbitofrontal cortex. Therefore, though we detected significant functional connectivity between the lateral orbitofrontal cortex and the striatum, it would be worth undertaking further work optimising signal within the orbitofrontal cortex to further investigate how it interacts with thalamostriatal connections during reversal learning.

In summary, we show that functional connectivity between the centromedian-parafascicular nuclei of the thalamus and the associative dorsal striatum contributes to adaptive behaviour. Functional connectivity is increased when processing negative versus positive outcomes and points to a simple system that utilises negative outcomes to detect potential changes in behavioural context and guide adaptive behaviour. This information may be used by the striatum to signal potential changes in context. Functional connectivity between the lateral orbitofrontal cortex and the associative dorsal striatum was also increased, but only during the stability phase. We believe this specificity is related to the role of the orbitofrontal cortex in flexibly implementing a change in behaviour when required. We also describe how task context might be represented and tracked within the prefrontal cortex. We suggest that activity within the frontal polar cortex is related to tracking the reliability of an alternative context to determine when a change in behaviour is required. Furthermore, we suggest that activity in the orbitofrontal cortex, anterior cingulate, insula, and inferior frontal gyrus may prepare neural architecture for change after receiving evidence that is incongruent with current expectations.

## Notes

### Competing Interest Statement

The authors have declared no competing interest.

